# No endospore formation confirmed in members of the phylum Proteobacteria

**DOI:** 10.1101/2020.07.23.219022

**Authors:** Polina Beskrovnaya, Doaa Fakih, Danielle L. Sexton, Shipei Xing, Mona Golmohammadzadeh, Isabelle Morneau, Dainelys Guadarrama Bello, Antonio Nanci, Tao Huan, Elitza I. Tocheva

## Abstract

Endospore formation is used by members of the phylum Firmicutes to withstand extreme environmental conditions. Several recent studies have documented endospore formation in species outside of Firmicutes, particularly in *Rhodobacter johrii* and *Serratia marcescens*, members of the phylum Proteobacteria. Here, we aimed to investigate endospore formation in these two species by using advanced imaging and analytical approaches. Examination of the phase-bright structures observed in *R. johrii* and *S. marcescens* using cryo-electron tomography failed to identify endospores or stages of endospore formation. We determined that the phase-bright objects in *R. johrii* cells were triacylglycerol storage granules and those in *S. marcescens* were aggregates of cellular debris. In addition, *R. johrii* and *S. marcescens* containing phase-bright objects do not possess phenotypic and genetic features of endospores, including enhanced resistance to heat, presence of dipicolinic acid, or the presence of many of the genes associated with endospore formation. Our results support the hypothesis that endospore formation is restricted to the phylum Firmicutes.

**Importance:** Endospore formation is a mechanism that allows bacteria to generate resilient dormant spores under harsh environmental conditions. Although this process has been traditionally restricted to the largely Gram-positive bacteria of the phylum Firmicutes, recent studies have also described endospores in some Proteobacteria. High complexity of endosporulation, reflected in extensive morphological transformations governed by hundreds of conserved genes, hinders its facile acquisition via horizontal gene transfer. Therefore, ability of distantly related bacteria to produce endospores would imply an ancient nature of this mechanism and potentially a pivotal role in species diversification and outer membrane biogenesis.

## Introduction

Spores represent a dormant state of bacteria that can persist for millions of years (1–3). Bacterial sporulation encompasses diverse modes, however, it is typically triggered by starvation and ultimately results in the production of metabolically inactive cells displaying increased resilience to stressors. For example, low nitrogen or carbon availability in Firmicutes can stimulate formation of endospores resistant to UV radiation, extreme pH, high temperature and pressure (4–6). Similarly, exosporulation in Actinobacteria and fruiting body production in *Myxococcus* have also been linked to nutrient limitation and can serve for preservation of genetic material under unfavourable environmental conditions (7–9). Despite the apparent similarities between these different types of sporulation, the underlying transformations are morphologically distinct and encoded by non-homologous pathways (10).

Endospore formation begins with asymmetric cell division, with the septum placed near one pole of the cell, and produces two cells with different fates (11–13). Upon septation, the smaller compartment becomes engulfed through a phagocytosis-like mechanism, yielding a prespore bound by two lipid membranes in the cytoplasm of the mother cell. Subsequent endospore maturation involves the synthesis of protective layers, such as the peptidoglycan-based cortex and proteinaceous coat. Metabolic inactivation is achieved by gradual dehydration of the core through replacement of water with dipicolinic acid (DPA) and calcium ions, and compaction of DNA with DNA-binding proteins. Together, these modifications account for the resistance properties of endospores (14). Ultimately, the spore is released upon lysis of the mother cell (15). In contrast, other modes of sporulation, such as those observed in Actinobacteria and *Myxococcus* sp., produce spores through morphological differentiation and cell division without engulfment.

Several studies in the past decade have reported, but not proven, formation of endospores in Proteobacteria (16, 17). While endosporulation has recently been confirmed in some Gram-negative bacteria, all of the identified organisms still belong to the phylum Firmicutes, highlighting the question of evolutionary origins of bacterial outer membrane (18). Additionally, sporulation involves tight cooperation of hundreds of genes distributed across the chromosome, hindering acquisition of this pathway through horizontal gene transfer (10, 19). Therefore, if confirmed, presence of the ability to form endospores across distantly related bacterial phyla suggests an ancient nature of the process and can provide clues to the characteristics of the last bacterial common ancestor (10). Here, we investigated two recent articles attributing sporulation to members of Proteobacteria (16, 17). Briefly, Girija *et al.* (17) described endospore production in the purple, non-sulfur bacterium *R. johrii*, strain JA192(T), a close relative of the model organism for bacterial photosynthesis *R. sphaeroides*. The second study by Ajithkumar *et al.* (16) reported endospore formation in *S. marcescens* subsp. *sakuensis* (strain no. 9; KRED^T^), a pathogenic bacterium that infects humans and causes bacteremia, urinary tract infection and wound infections (20). Thus, confirmation and further characterization of endospore formation in these organisms can bring valuable insight into the physiology of these species and the role of endospore formation in diversification and speciation of modern phyla.

In this study, we employed cutting edge structural biology techniques, such as cryo-electron tomography (cryo-ET), correlative light and electron microscopy (CLEM), and energy-dispersive X-ray spectroscopy (EDX), as well as biochemical and microbiological approaches, to characterize endospore formation in *R. johrii* and *S. marcescens*. Our results showed that *R. johrii* and *S. marcescens* were unable to form endospores as previously reported (16, 17). Further analyses indicated that the putative spores in *R. johrii* were lipid storage granules rich in triacyclglycerols (TAGs), and the phase-bright objects in *S. marcescens* were aggregates of cellular debris. Overall, our observations contradict the previously published studies by Girija *et al.* and Ajithkumar *et al.*, and support the observation that these members of Proteobacteria are unable to form endospores.

## Materials and Methods

### Bacterial strains and growth conditions

*R. johrii* and *S. marcescens* cells were purchased from Leibniz-Institut DSMZ bacterial strain collection. *R. johrii* JA192 cells (DSMZ 18678) were cultivated as previously described by Girija *et al.* (17). Briefly, cells were grown aerobically at room temperature in *R. sphaeroides* solid and liquid medium comprising 4 mM KH_2_PO_4_, 1 mM MgCl_2_·6H_2_O, 7 mM NaCl, 22 mM NH_4_Cl, 0.04 mM CaCl_2_·2H_2_O, 17 mM sorbitol, 28 mM sodium pyruvate, 1.5 mM yeast extract, 1 L distilled water, pH 7.0, 1 ml trace element solution SL7, and 20 ng Vitamin B_12_ solution for 2 days for vegetative cells or 7 days to induce production of phase-bright objects. Additionally, cells were grown in Luria-Bertani (LB) broth at 30 °C with agitation for 2 days, and either harvested as the vegetative growth control or subsequently inoculated 1:100 into modified M9 medium for an additional 7 days to induce formation of phase-bright objects. The modified M9 medium contained 47.8 mM Na_2_HPO_4_, 22 mM KH_2_PO_4_, 8.56 mM NaCl, 18.7 mM (3.74 mM for limited nitrogen) NH_4_Cl, 1 mM MgSO_4_, 0.3 mM CaCl_2_, 0.4 % (w/v) glucose, 1 μg/L biotin, 1 μg/L thiamine, 31 μM FeCl_3_·6H_2_O, 12.5 μM ZnCl_2_, 2.5 μM CuCl_2_·2H_2_O, 2.5 μM CoCl_2_·2H_2_O, 5 μM MnCl_2_·4H_2_O, 2.5 μM Na_2_MoO_4_·2H_2_O. *S. marcescens* cells (DSMZ 30121) were cultivated in LB broth at 32 °C with shaking at 200 rpm, as previously described by Ajithkumar *et al.* (16), for 7 days for vegetative growth or 65 days to induce formation of phase-bright objects. *B. subtilis* strain PY79 was chosen as the positive control for endospore formation, and cells were cultivated in LB broth at 37 °C with shaking at 200 rpm overnight for vegetative growth or for 3 days to induce sporulation. LB agar was used for cultivation on plates for *S. marcescens* and *B. subtilis*.

### Detection of phase-bright objects using phase-contrast light microscopy

*R. johrii* and *S. marcescens* cultures were pelleted and washed with 1x phosphate buffer saline (PBS, pH 7.4) composed of 137 mM NaCl, 27mM KCl, 10 mM Na_2_HPO_4_, 1.8 mM KH_2_PO_4_. Cells were imaged with an upright Zeiss Axio Imager M2 Microscope (Carl Zeiss, Oberkochen, Germany) equipped with a 506 monochrome camera, and a 100x oil objective lens with a numerical aperture (NA) of 1.46.

### Sample preparation for correlative light and cryo electron tomography

*R. johrii* cells were lightly fixed using 2.5 % paraformaldehyde in 30mM phosphate buffer for 15 min, washed twice and resuspended in 150 mM phosphate buffer. Bacterial cells were loaded onto Cu Finder R 2/2 EM grids (Electron Microscopy Sciences, Hatfield, USA), coated with 1 mg/ml poly-L-lysine, and subsequently imaged at room temperature as described above. Following room temperature light microscopy, 20-nm colloidal gold particles (UMC Utrecht, Netherlands) were added and samples were plunge-frozen into liquid ethane-propane mix cooled at liquid nitrogen temperatures with a Mark IV Vitrobot (Thermo Fisher Scientific), maintained at room temperature and 70 % humidity. Cryo-ET was conducted on cells with phase-bright signal as described below.

### Cryo-ET sample preparation

For standalone cryo-ET experiments, samples were mixed with 20-nm colloidal gold particles, loaded onto glow-discharged carbon grids (R2/2, Quantifoil) and plunge-frozen into liquid ethane-propane mix cooled at liquid nitrogen temperatures with a Mark IV Vitrobot(Thermo Fisher Scientific) maintained at room temperature and 70 % humidity.

### Cryo-ET data collection

For both standalone cryo-ET and CLEM experiments, tilt-series were collected on an 300kV Titan Krios transmission electron microscope (Thermo Fisher Scientific) equipped with a Falcon 2 camera. Tilt series were collected at 14-18K x nominal magnification, 1-3 degrees oscillations and a final dose of 30-150 e^−^/A^2^. Three-dimensional reconstructions were calculated with IMOD using the weighted back projection method (21).

### Correlative LM and SEM with EDX analysis

*R. johrii* cells were fixed by 12.5% paraformaldehyde in 150 mM sodium phosphate buffer (73.6 mM K_2_HPO_4_, 26.4 mM KH_2_PO_4_) pH 7.5, then washed three times with 150mM sodium phosphate buffer (22). Glow-discharged Cu R2/2 grids were coated with poly-L-lysine hydrobromide solution (1 mg/ml) and dried for 30 mins at 60 °C. Fixed cells were loaded onto the grids and immediately imaged with LM in 1x PBS to identify phase-bright objects. The grids were subsequently air dried, and regions of interest identified with LM were examined with SEM using JEOL JSM-7400F (JEOL Ltd., Tokyo, Japan) operated at 5 kV without any coating. EDX analysis with a silicon-drift detector (Octane, EDAX Inc., Mahwah, NJ, USA) at 10 kV was used for semi-quantitative elemental analysis of regions of interest.

### Whole-cell lipidomic analysis of R. johrii

*R. johrii* cells displaying phase-bright properties were cultivated as described above, harvested by centrifugation (20 min at 5,000 rpm) and washed twice in sterile H_2_O. Cell pellets were then lyophilized overnight and stored at room temperature for up to 7 days. *R. johrii* cultures grown in LB for 2 days and lacking phase-bright objects were chosen as the negative control. For the whole-cell lipidomics analysis, methyl tert-butyl ether (MTBE)-based membrane lipid extraction protocol was used with modifications (23). Briefly, samples in 1.5 ml Eppendorf vials were first mixed with 300 μl ice-cold methanol and 10 μl internal standards. The mixture was then sonicated in ice-water bath for 15 min for protein precipitation. 1 ml MTBE was added to the mixture, followed by vortex mixing for 20 min at room temperature for thorough lipid extraction. Next, 200 μl LC-MS grade water was added to induce phase separation, and the samples were further mixed for 30 s. After settling for 10 min, the upper layer, containing the lipids, was transferred to new Eppendorf vials. To dry the lipid samples, the solvent was evaporated using a vacuum concentrator at 4°C. 100 μL isopropanol/acetonitrile (1:1, v/v) was added to reconstitute the dried residue. The reconstituted solution was vortexed for 30 s and centrifuged at 14,000 rpm at 4°C for 15 min. The resulting supernatants were transferred to glass inserts for liquid chromatography-tandem mass spectrometry (LC-MS/MS) analysis. Only lipids above the noise level (1000 average intensity) were considered in the analysis. A cut off value of at least 2x increase in average intensity, and the p-value threshold of 0.01 was used to determine significant increase in lipid species.

### Heat inactivation and counting of endospores

After induction of the phase-bright objects in *R. johrii, S. marcescens*, and *B. subtilis* as described above, cells were washed with sterile, deionized water, spun at 10,000 g for 10 min, and resuspended in chilled water. Suspensions of *R. johrii* and *B. subtilis* were heated to 80 °C, and *S. marcescens* to 60 °C, for 15 min, 30 min and 1 hr, as described previously (16, 17). After the heat treatment, the samples were centrifuged for 10 min at 10,000 g. The pellets were washed five times to remove cellular debris, then plated on solid media. *R. johrii* was incubated at 30 °C for 7 days, *S. marcescens* was incubated at 32 °C for 7 days, and *B. subtilis* was incubated at 37 °C overnight, and plates were subsequently examined for viable growth.

### Dipicolinic acid (DPA) detection

Following the detection of phase-bright objects, DPA was detected as previously described (24). Briefly, cultures of *R. johrii*, *S. marcescens*, and *B. subtilis* containing ~ 10 mg dry weight were autoclaved for 15 minutes at 15 lb/in^2^. The suspensions were cooled to ambient temperature, acidified with 0.1 ml of 1.0 N acetic acid and incubated for 1 hour to cluster the insoluble material. To remove cellular debris, the suspensions were centrifuged at 1,500 g for 10 minutes. To each 4 ml of supernatant, 1 ml of 1 % Fe(NH_4_)_2_(SO_4_)_2_·6H_2_0 and 1 % ascorbic acid in 0.5 M acetate buffer (pH 5.5) was added. Colorimetric shift at 440 nm was compared to a standard curve prepared with pure DPA (Sigma-Aldrich, Oakville, Canada).

### Detection of genes required for endospore production

List of conserved genes, previously identified as essential for endospore formation, was compiled based on the studies by Galperin *et al.* and Meeske *et al.* (5, 25). The respective amino acid sequences, encoded by the model strain *B. subtilis* 168, were then retrieved from the UniProt database (https://www.uniprot.org/). Sequence searches were performed against the genomes of *R. johrii* JA192 and *S. marcescens* subsp. *sakuensis* KRED^T^, the two strains originally described by Girija *et al.* and Ajithkumar *et al.*, respectively (16, 17), using tBLASTn (https://blast.ncbi.nlm.nih.gov/Blast.cgi). Positive hits were defined by a BLAST score >60 and sequence identity >30 %.

## Results and Discussion

### Prolonged incubation induces formation of phase-bright objects in *R. johrii* and *S. marcescens*

For initial assessment of the previously reported endospore formation, we cultivated *R. johrii* and *S. marcescens* according to the published conditions and examined the cultures with phase-contrast LM (Fig. 1). Vegetative *R. johrii* cells appeared phase-dark (Fig. 1A), however, upon 7-day incubation phase-bright objects were observed either at one pole or mid-cell (Fig. 1B, black arrows). Vegetative *S. marcescens* cells also appeared phase-dark (Fig. 1C). Although phase-bright objects were occasionally visible at mid-cell following 65-day incubation (Fig. 1D, black arrows), the majority of culture was dead and appeared as “ghost” cells (Fig. 1D, white arrows). Altogether, our results recapitulate reports of formation of phase-bright objects in *R. johrii* and *S. marcescens* following extended incubation in nutrient-limited conditions.

**Figure 1.**
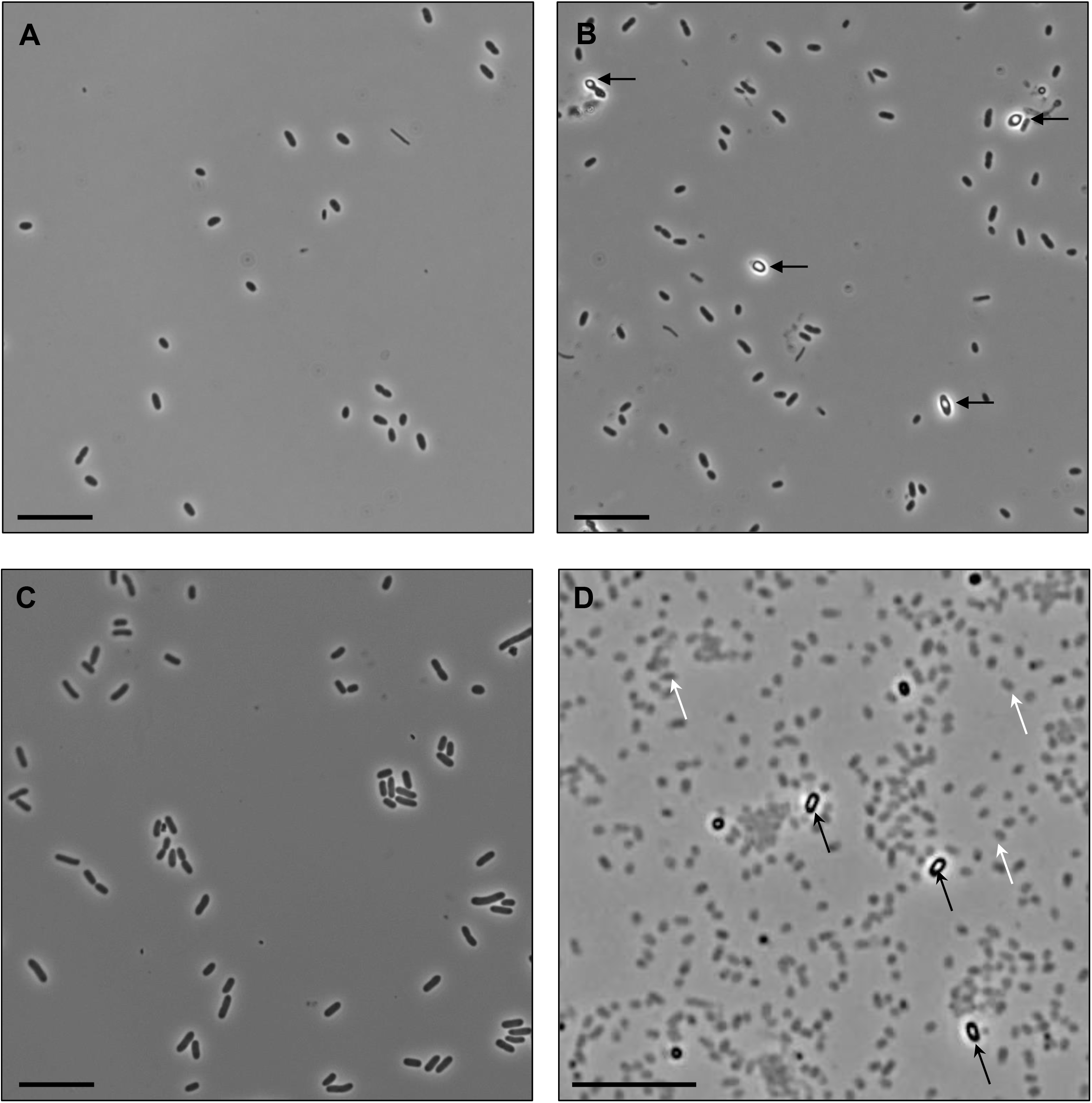
Phase-contrast light microscopy of *R. johrii* and *S. marcescens* cells. A) 2-day old *R. johrii* cells lack phase-bright objects; B) 7-day old *R. johrii* cells with phase-bright objects (black arrows). C) 7-day old *S. marcescens* cells lack phase-bright objects. D) After 65 days, *S. marcescens* cells show two kinds of cell morphologies: phase-bright (black arrows), and “ghost” cells (white arrows). Scale bar 10 μm.

### Characterization of phase-bright objects in *R. johrii*

To further characterize the phase-bright objects observed in *R. johrii*, we performed correlative LM and cryo-ET experiments on phase-bright and phase-dark cells following extended incubation (Fig. 2). Tomograms of *R. johrii* cells with phase-bright objects revealed the presence of intracellular granules, which were highly sensitive to the electron beam, as represented by the sample damage (Fig. 2A). Beam sensitivity was detected regardless of the total dose used (25-150 e^−^/A^2^), suggesting that the granules were rich in lipids. Further, the spherical nature of the granules resembled previously characterized storage granules (SG) in bacterial cells (26). No evidence of sporulation-associated morphological changes, such as engulfing membranes, presence of immature or mature spores in the sample (n=40), was observed, indicating that the phase-bright objects were not endospores. Finally, cells with phase-bright objects always displayed 1-3 of the 100-250 nm diameter granules (Fig. 2A), whereas the phase-dark cells lacked the presence of granules (Fig. 2B). Thus, our observations suggest that the phase-bright objects were likely lipid-containing SGs.

**Figure 2.**
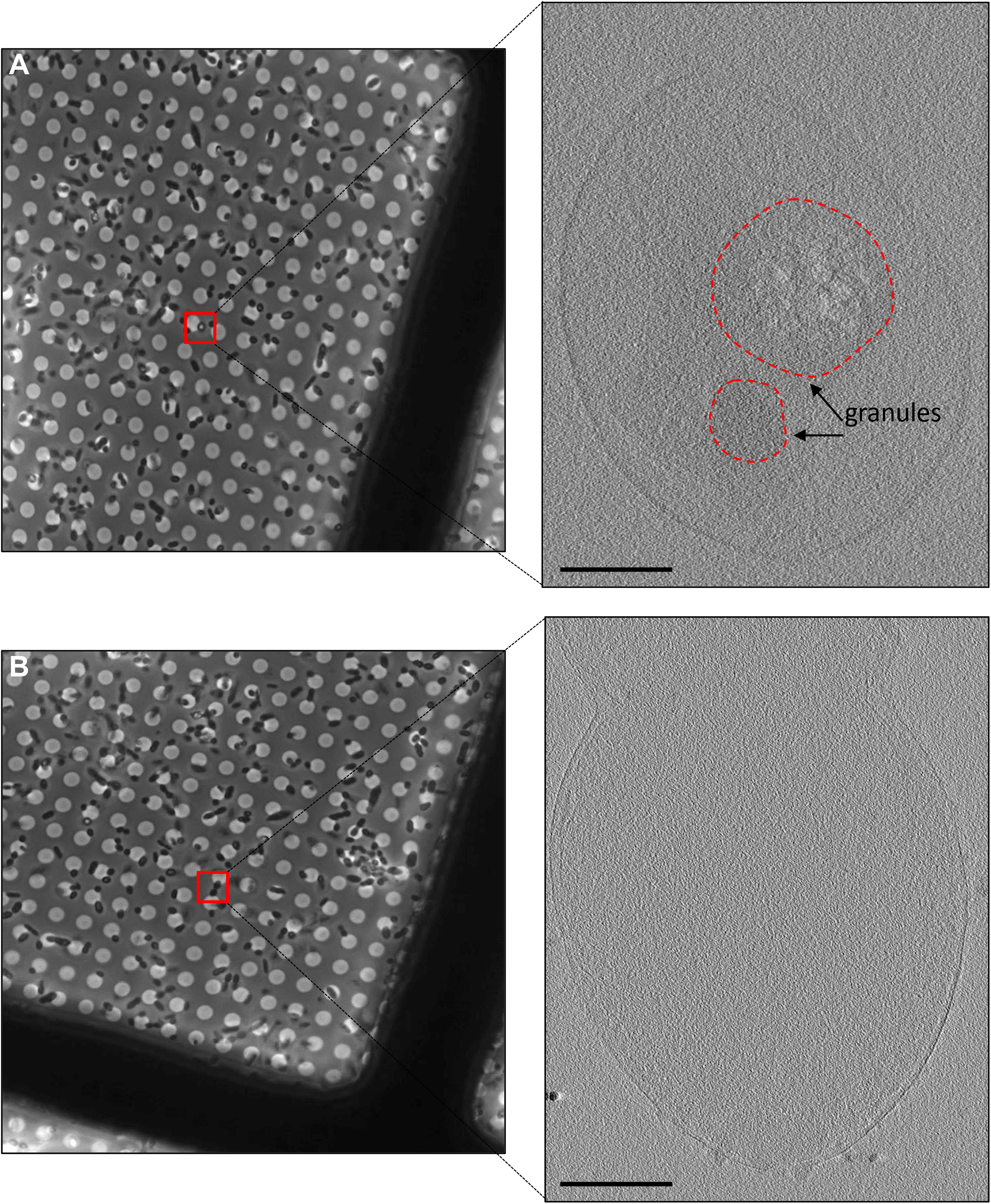
Correlative light and cryo-ET of *R. johrii.* (A) Left: Phase-contrast microscopy image of a *R. johrii* cell (boxed) displaying a phase-bright object. Right: tomographic slice of the same cell showing two granular structures. (B) Left: Phase-contrast microscopy image of *R. johrii* cells (boxed) lacking phase bright objects. Right: tomographic slice of the same cell showing lack of sub-cellular structures. Scale bar 200nm.

To determine the composition of the storage granules, we performed correlative LM and SEM in combination with EDX compositional analysis. Cells possessing the putative storage granules were identified with phase-contrast microscopy (Fig. 3A) and examined in higher resolution with SEM (Fig. 3B). Correlative LM and SEM was then used to guide EDX analysis, so that spectra were collected from a region containing the putative storage granules and a cytoplasmic region lacking the storage granules (Fig. 3C). Elemental analysis of the storage granule (blue spectrum) revealed counts for carbon (C) 80.24 %, oxygen (O) 13.26%, and copper (Cu) 6.5% (due to the copper of the EM grid). Cytoplasmic analysis (red spectrum) revealed lower counts for carbon (C) 61.7% and oxygen (O) 10.82%, copper (Cu) at 6.59%, and elevated counts for nitrogen (N) 20.9% (Fig. 4C).

**Figure 3.**
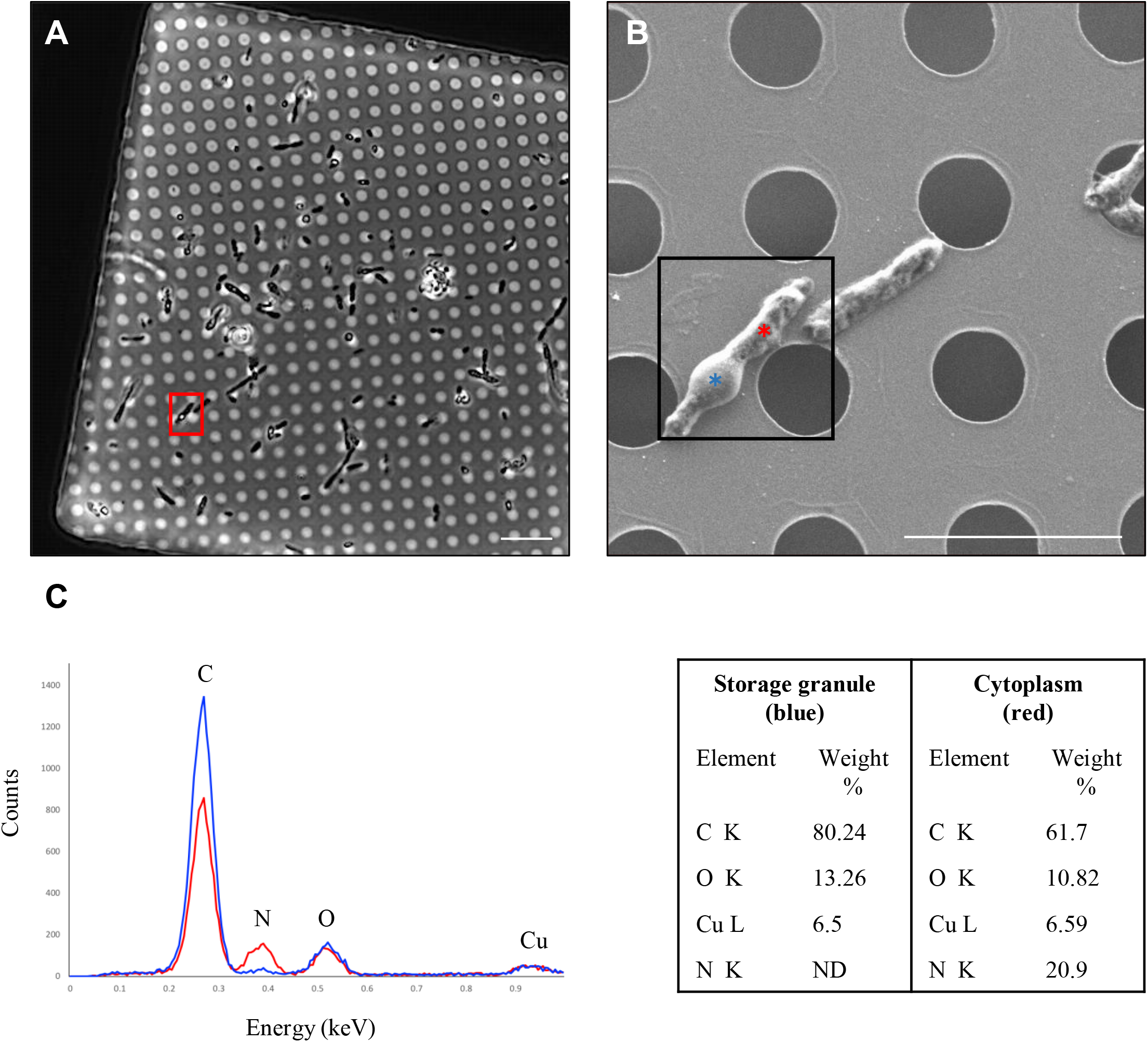
Correlative LM and SEM of *R. johrii* for storage granule characterization with EDX. (A) LM image of *R. johrii* shows the presence of storage granules (phase-bright objects) inside a cell (red square). (B) The same cell as in panel A imaged with SEM. Areas corresponding to the storage granule and cytoplasm are depicted as blue and red stars, respectively. (C) Elemental composition of the storage granule (blue) and cytoplasm (red) using EDX semi-quantitative analysis. Major peaks are assigned and data is summarized in a table format. Scale bar 10 μm (A), 5 μm (B). ND – non-detected.

**Figure 4.**
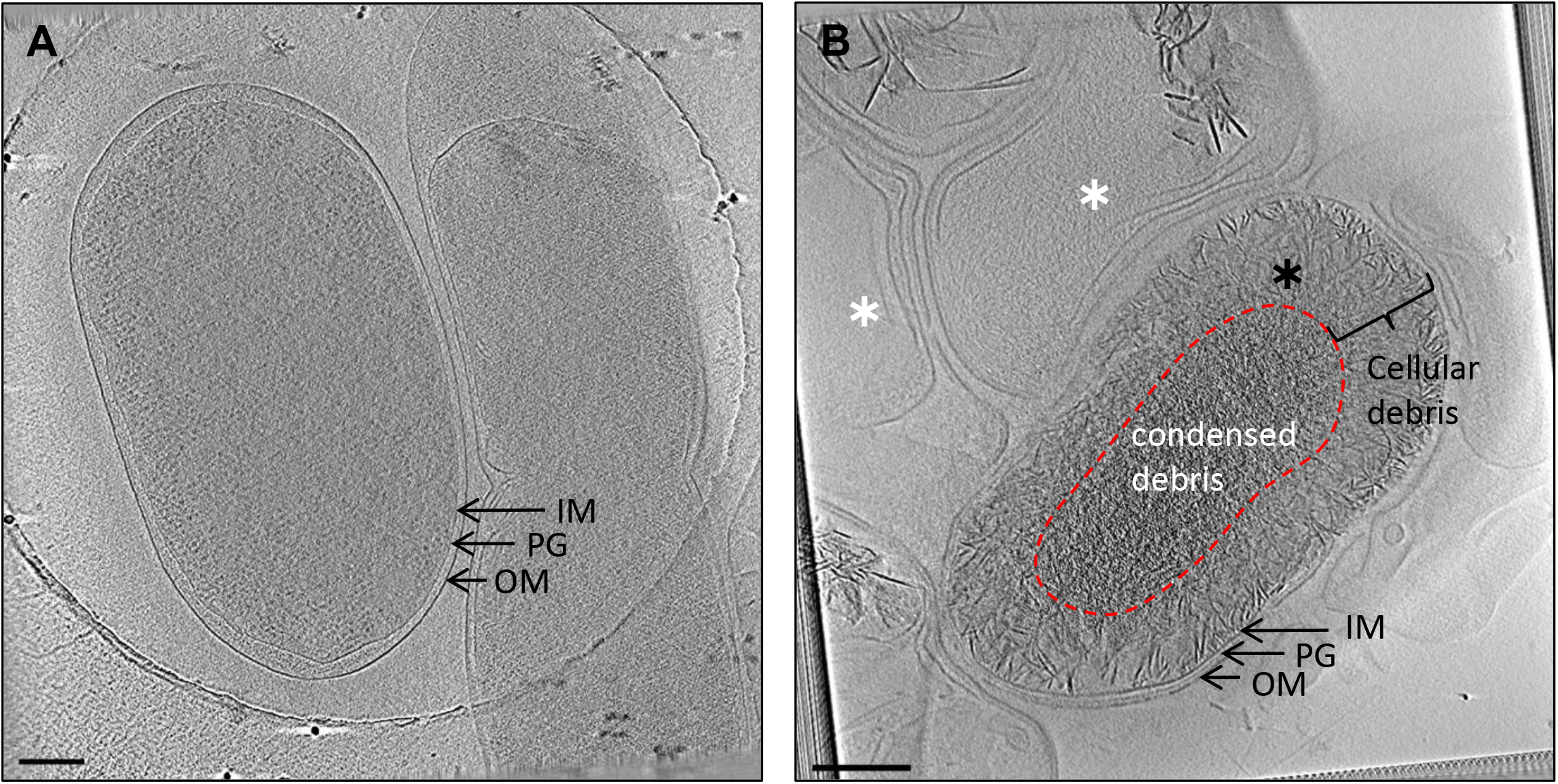
Cryo-ET of *S. marcescens*. Tomographic slices through: (A) Vegetative cells from a 2-day old culture; (B) Cells from a 65-day old culture showing phase-bright objects. Panel B shows two cell types: cells with accumulated cellular debris (black star) and “ghost cells” void of cellular material (white star). Scale bar 200 nm. IM, inner membrane; PG, peptidoglycan; OM, outer membrane.

Based on the cryo-ET and EDX data, we hypothesized that the granules observed in *R. johrii* were composed of lipids, as lipids are enriched in carbon and oxygen atoms. To characterize the nature of the granular composition, we performed whole-cell lipidomics analysis of *R. johrii* 7-day old culture expressing phase-bright objects (granules) against *R. johrii* cells grown for 2 days and lacking phase-bright objects as the negative control. Cells producing putative storage granules where enriched in several lipids, the most abundant of which were triacylglycerols (TAGs) and phosphatidylethanolamines (PEs) (Table 1). Because PEs are typical membrane lipids, the increased levels observed under starvation conditions suggested that cells remodel their membrane composition to account for the environmental changes. TAGs are nonpolar triacylglycerols that occur as insoluble inclusions in bacteria and are considered a major source of energy (27, 28). TAGs have been shown to accumulate in actinobacteria and mycobacteria as either peripheral deposits associated with the cell envelope, or as inclusion bodies in the cytoplasm (29). Previously, *in vitro* studies showed that mycobacteria accumulated TAG and wax ester when subjected to stresses, such as low oxygen, high CO_2_, low nutrients and low pH (29–31). Similarly, we observed an increased propensity to form the phase-bright objects in *R. johrii* cells incubated under low-nitrogen conditions in defined media. Therefore, it is likely that *R. johrii* utilizes TAG storage as an adaptive strategy in response to starvation, allowing cells to enter stationary phase and survive for longer periods of time. We thus conclude that the granules observed as phase-bright objects in *R. johrii* were storage granules enriched in TAGs.

**Table 1.** Lipidomics analysis of whole *R. johrii* cells.

| Lipid<sup>a</sup> | Lipid class | Fold change<br>R.j + / R.j - | P-value |
| --- | --- | --- | --- |
| PE 33:1; PE 16:0-17:1<sup>c</sup> | PE | 147.84 | 4.71E-09 |
| TAG 58:1; TAG 16:0-24:0-18:1 | TAG | 93.81 | 1.81E-08 |
| TAG 52:3; TAG 16:0-18:1-18:2 | TAG | 67.19 | 1.24E-08 |
| TAG 54:5; TAG 18:1-18:2-18:2 | TAG | 60.41 | 1.37E-06 |
| TAG 52:2; TAG 18:0-16:1-18:1 | TAG | 57.72 | 1.93E-07 |
| TAG 54:4; TAG 18:1-18:1-18:2 | TAG | 54.46 | 2.37E-08 |
| TAG 52:1; TAG 16:0-18:0-18:1 | TAG | 49.15 | 1.22E-10 |
| TAG 56:2; TAG 16:0-18:1-22:1 | TAG | 48.89 | 5.37E-09 |
| TAG 50:1; TAG 16:0-16:0-18:1 | TAG | 46.43 | 7.39E-09 |
| TAG 54:2; TAG 18:0-18:1-18:1 | TAG | 44.82 | 2.37E-09 |
| TAG 58:2; TAG 16:0-18:1-24:1 | TAG | 42.04 | 9.59E-09 |
| TAG 52:2; TAG 16:0-18:1-18:1 | TAG | 41.06 | 5.93E-09 |
| TAG 56:1; TAG 16:0-22:0-18:1 | TAG | 39.72 | 8.65E-10 |
| TAG 54:1; TAG 18:0-18:0-18:1 | TAG | 39.35 | 1.15E-08 |
| TAG 54:3; TAG 18:0-18:1-18:2 | TAG | 17.15 | 3.93E-08 |
| TAG 50:2; TAG 16:0-16:1-18:1 | TAG | 12.30 | 1.90E-08 |
| PE 32:0; PE 16:0-16:0 | PE | 11.55 | 1.73E-06 |
| PC 39:3 | PC | 10.77 | 1.06E-08 |
| PE 32:1; PE 16:0-16:1 | PE | 7.68 | 3.84E-07 |
| TAG 48:1; TAG 14:0-16:0-18:1 | TAG | 4.94 | 2.00E-05 |
| PE 35:2; PE 17:1-18:1 | PE | 3.09 | 6.92E-06 |
| PC 36:4 | PC | 3.03 | 4.93E-06 |
| TAG 48:1; TAG 16:0-16:0-16:1 | TAG | 2.52 | 4.51E-03 |
| PC 32:1 | PC | 2.18 | 4.46E-07 |
| PC 34:1; PC 16:0-18:1 | PC | 2.15 | 6.50E-08 |
| DAG 36:2; DAG 18:1-18:1 | DAG | 2.08 | 5.34E-08 |
| PC 34:2; PC 16:1-18:1 | PC | 2.02 | 1.57E-06 |

<sup>a</sup> Abbreviations: phosphatidyl ethanolamine, PE; triacylglycerols, TAG; phosphatidylcholine, PC; diglyceride, DAG
<sup>b</sup> the total lipid composition of *R. johrii* expressing storage granules (R.j +) was compared to a fresh *R. johrii* culture lacking storage granules (R.j -)
<sup>c</sup> PE is a lipid class with two acyl chains (R1 and R2 in B). PE 16:0-17:1 indicates that the chain lengths are 16 carbons and 17 carbons, and the saturation degrees are 0 and 1, respectively. PE 33:1 is the simpler form of PE 16:0-17:1.

### Characterization of phase-bright objects in *S. marcescens*

Tomograms of *S. marcescens* were collected on 2-day and 65-day old cultures (Fig. 4). At 2 days, we observed regular morphology of vegetative cells, displaying cell envelope architecture typical for Gram-negative bacteria (Fig. 4A). *S. marcescens* grown for 65 days revealed the presence of two kinds of morphologies: cells packed with cellular debris (black star), and cells void of any cellular material (white star) (Fig. 4B), likely correlating to cells containing phase-bright objects and “ghost” cells identified using LM, respectively. Extensive survey of the sample (n=80) did not reveal any cells possessing intracellular membranes, or morphologies suggestive of engulfing membranes or stages of sporulation. Neither of the two identified morphologies displayed any features similar to a cortex or proteinaceous spore coat characteristic of mature endospores. Additionally, we did not observe accumulation of storage granules within cells. Together, these results suggest that the appearance of phase-bright objects in *S. marcescens* was the result of accumulation of cellular debris, and dehydration.

### Proteobacteria do not possess features of endospores following extended incubation

Independent of imaging-based methods, endospores have traditionally been identified in samples through heat resistance and increased concentration of intracellular DPA. To verify the results of our cryo-ET experiments, we first investigated the heat resistance properties of *R. johrii* and *S. marcescens* following prolonged incubation and subsequent exposure to high temperatures. Despite an extended recovery period of 7 days, no viable *R. johrii* cells were observed on solid media following 15, 30 and 60 min incubation at 80 °C. Similarly, viable cells were not isolated from *S. marcescens* cultures incubated at 60 °C for 15, 30, and 60 min. In contrast, *B. subtilis* cultures producing endospores and treated at 80 °C for 15, 30, and 60 min yielded viable growth on solid media after a 24 hr recovery period. Thus, we were unable to replicate the results of Girija *et al.* and Ajithkumar *et al.*, who found viable cells following heat treatment of *R. johrii* at 80 °C for 20 min, and *S. marcescens* at 60 °C for 15 min, respectively. Additionally, we quantitatively analyzed the presence of DPA in cultures of *R. johrii* and *S. marcescens* displaying phase-bright objects using a colorimetric method. Whereas the purified endospores of *B. subtilis* contained 6.74 μg/ml DPA, no detectable amounts of DPA were observed in *R. johrii* and *S. marcescens* after prolonged cultivation. Collectively, these results indicate that *R. johrii* and *S. marcescens* cells do not possess the classic phenotypic features that are associated with endospore formation.

### Minimal subset of genes required for endospore formation not conserved in the Proteobacteria

Endospore formation relies on expression of hundreds of genes in a highly regulated manner (19, 32, 33). For example, over 500 genes have been previously implicated in sporulation in the model firmicute *B. subtilis* (32). However, establishment of the minimal subset of genes required for endospore formation remain elusive, as many of the identified targets carry out redundant functions, *e.g.* histidine kinases, or are part of general pathways loosely associated with sporulation, such as iron uptake and DNA repair proteins (34). Consistently, several homologs to genes linked to sporulation have been detected in other phyla, including Proteobacteria, but have been shown to play regulatory roles in distinct processes, such as cell division and development (35, 36). Hence, possession of genes annotated as sporulative should not be considered concrete evidence to support sporulation capacity in a given species (5). Nevertheless, we investigated the genomes of *R. johrii* and *S. marcescens* for presence of genes that are conserved among all spore-forming bacilli and clostridia, and have been shown to play pivotal roles in endospore formation through functional studies (Table 2–3) (5, 25). Our analysis showed that that both *R. johrii* and *S. marcescens* completely lack the SpoIIDMP peptidoglycan remodeling complex required for spore cortex formation, the SpoIIQ-SpoIIIAA-AH channel complex involved in communication between the mother cell and the prespore and facilitating regulation of endospore maturation, as well as the major protein coat assembly components, such as SpoIVA and Alr (12, 14, 37). Further, the master regulator of sporulation encoded by all endospore-formers, Spo0A, is absent in *R. johrii*. Both strains also lack homologs to three out of four sporulation sigma factors, SigF, SigE and SigK. Finally, *R. johrii* and *S. marcescens* do not possess DapB, required for production of dipicolinic acid which plays a major role in dehydration of the spore core and, therefore, resistance and dormancy (14). Therefore, our analysis confirms the lack of those conserved genes in the genomes of *R. johrii* and *S. marcescens*. In addition, Ajithkumar *et al.* was also unable to detect genes related to endospore formation in *S. marcescens* (16).

**Table 2.**
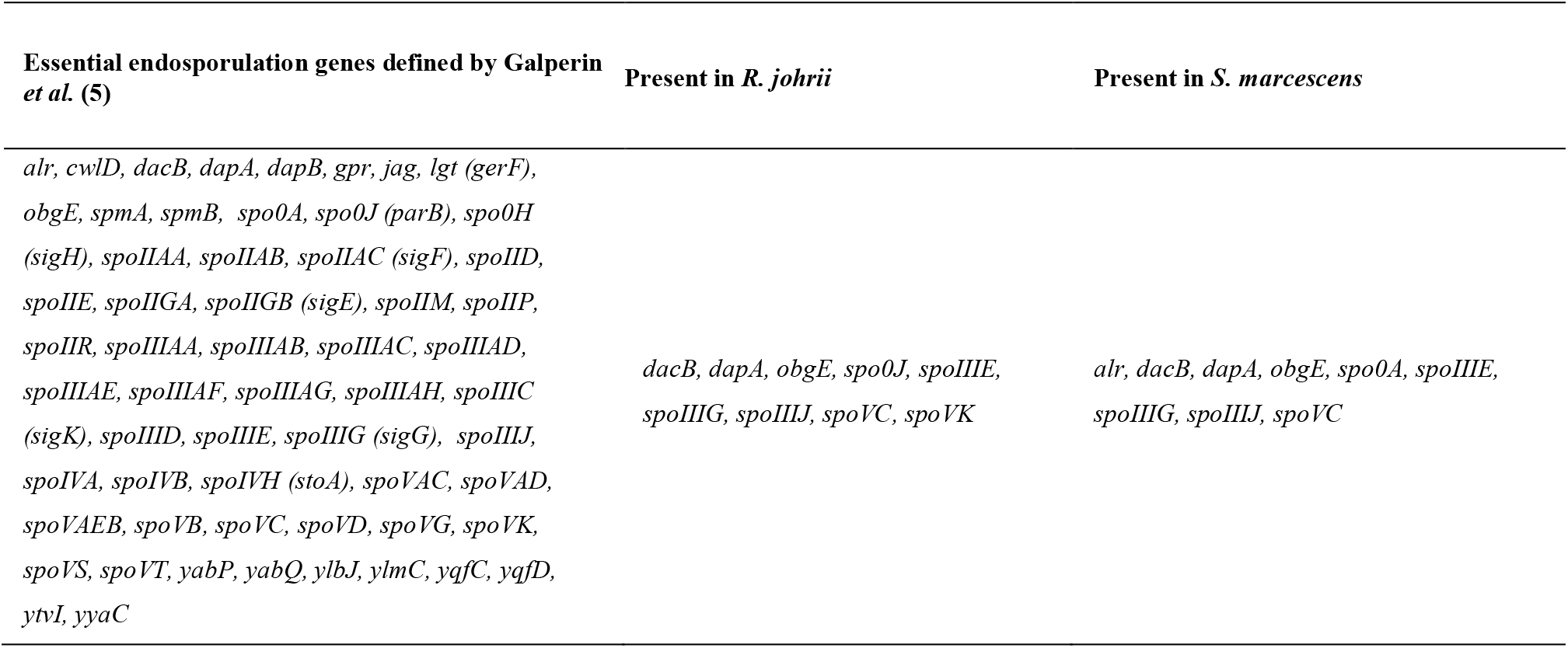
Analysis for presence of the minimal subset of endosporulation genes in *R. johrii* and *S. marcescens*.

**Table 3.**
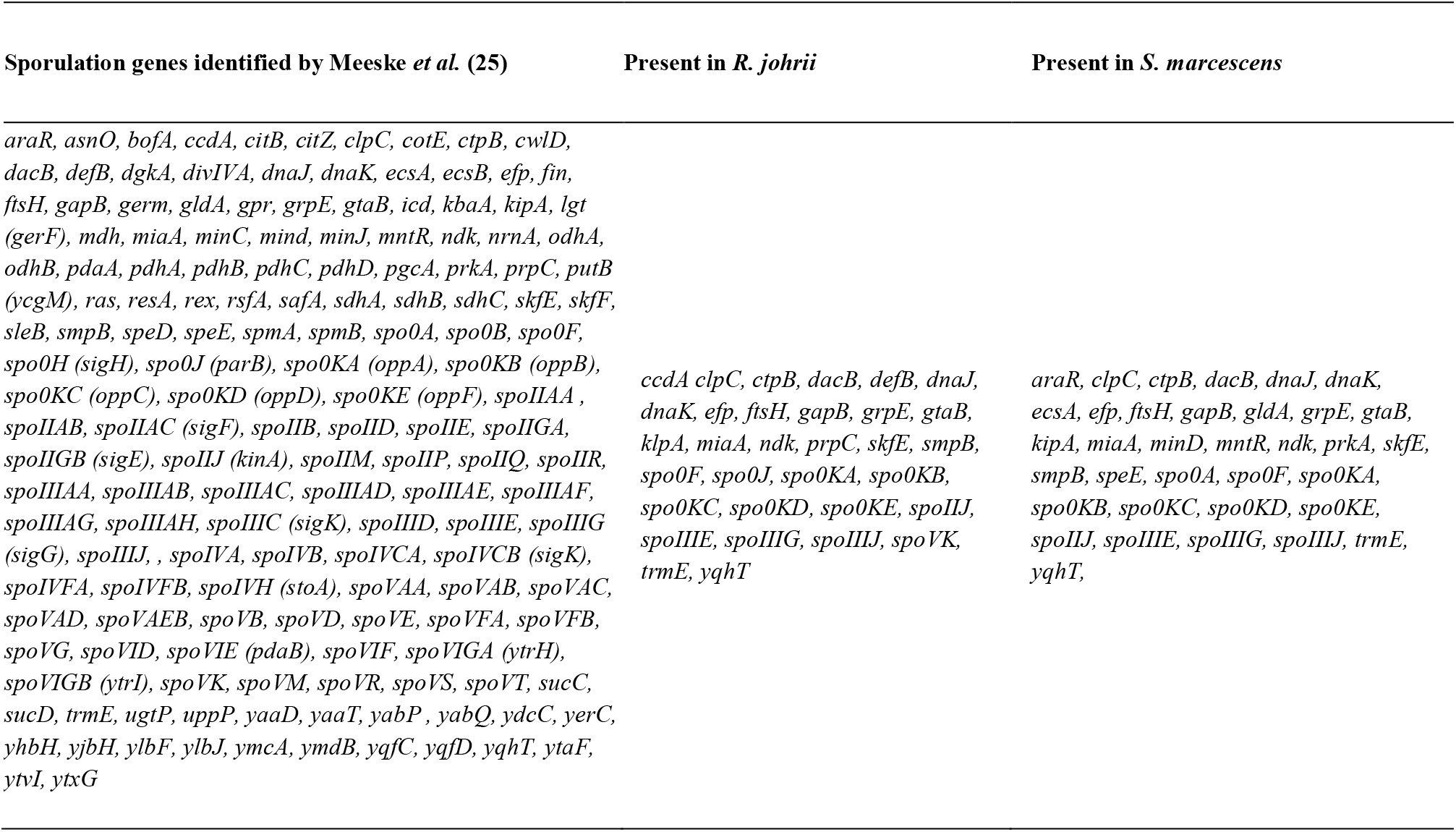
Survey for presence of additional endosporulation genes in *R. johrii* and *S. marcescens*.

### Concluding remarks

Although endospore formation is considered a hallmark of the Firmicutes phylum (4, 5, 38), endospore production had been reported outside of Firmicutes, particularly in two members of the phylum Proteobacteria (16, 17). These findings may affect our understanding of the evolutionary events surrounding outer membrane biogenesis and the significance of endospore formation in cell differentiation. Here, using cutting-edge microscopy techniques, and biochemical, microbiological, as well as bioinformatics approaches, we showed that the phase-bright objects observed in *R. johrii* and *S. marcescens*, are storage granules and cellular debris, respectively. We did not observe mature spores or stages of endospore formation *in vivo*, and failed to detect the pivotal biochemical and genomic features of endospore-producing bacteria in these organisms. Our findings thus demonstrate that *R. johrii* and *S. marcescens* are unable to form true endospores, which is in contrast to the results described by Girija *et al.* (17) and Ajithkumar *et al.*(16). Since we used the most-advanced imaging techniques currently available to study whole-cell bacteria and their ultrastructure, previous results could be due to the presence of contamination with spore-forming bacteria or misinterpretation of methodology artifacts.

## Acknowledgements

We thank Dr. Kaustuv Basu at the Facility for Electron Microscopy Research (FEMR) of McGill University and Dr. Claire Atkinson at the High Resolution Macromolecular Cryo-Electron Microscopy Facility at the University of British Columbia for help with microscope operation and data collection. Work in the EIT lab was supported by Natural Sciences and Engineering Research Council of Canada Discovery Grant (RGPIN 04345) and CRC Tier 2.

